# Preserved Embedding of Neural Population Activity for Brain Computer Interfaces

**DOI:** 10.64898/2026.02.05.704020

**Authors:** Hongru Jiang, Zhe Liu, Xiangdong Bu, Antong Liu, Xingyu Ji, Yao Chen

## Abstract

Inferring neural population dynamics from motor cortex recordings is essential for brain-computer interface (BCI) control. However, neural activity patterns during similar behaviors exhibit systematic variability across recording sessions, subjects, and task contexts due to neural plasticity, anatomical differences, and varying movement distances. This variability degrades BCI decoder performanc when applied to different sessions, subjects, and task contexts. Here, we introduce Multi-Aligned Neural Data Transformer (MANDT) that extracts consistent embeddings of the motor cortical dynamics. By learning these consistent embeddings, MANDT enables generalization of BCI decoding: a decoder trained on data from one session maintains robust performance when applied to new sessions without recalibration. We validated our approach on neural recordings from rhesus monkeys performing the delayed-reach task. Lastly, we show that CEBRA can be used for the robotic control, enabling direct neural control of a humanoid robot, allowing monkeys to guide complex robotic manipulation tasks through brain signal.

## 1 Introduction

Brain-computer interfaces (BCIs) restore motor function to paralyzed patients by decoding neural activity from the motor cortex into prosthetic movements [1–5]. Modeling the relationship between neural activity and corresponding behavior is critical for developing high-performance BCIs. Various approaches, including Wiener filter [6], Kalman filters [2], optimal linear estimator [1], and long short-term memory networks [7] have been employed for behavioral decoding. However, neural recordings are are highly variable across contexts, limiting decoder generalizability [8] (Fig. 2a, b). Specifically, three sources of variability pose challenges: First, session-to-session variability occurs as extracellular environment changes (e.g., neuron death or electrode drift [9, 10]) across days. Second, subject-to-subject variability arises from differences in brain anatomy and electrode placement across individuals. Third, task-to-task variability exits due to differences in task parameters (e.g., target distance, or spatial dimensions of movement). The variability across session, subject and task degrade BCI performance, leading to extensive and time-consuming recalibration.

**Fig. 2:**
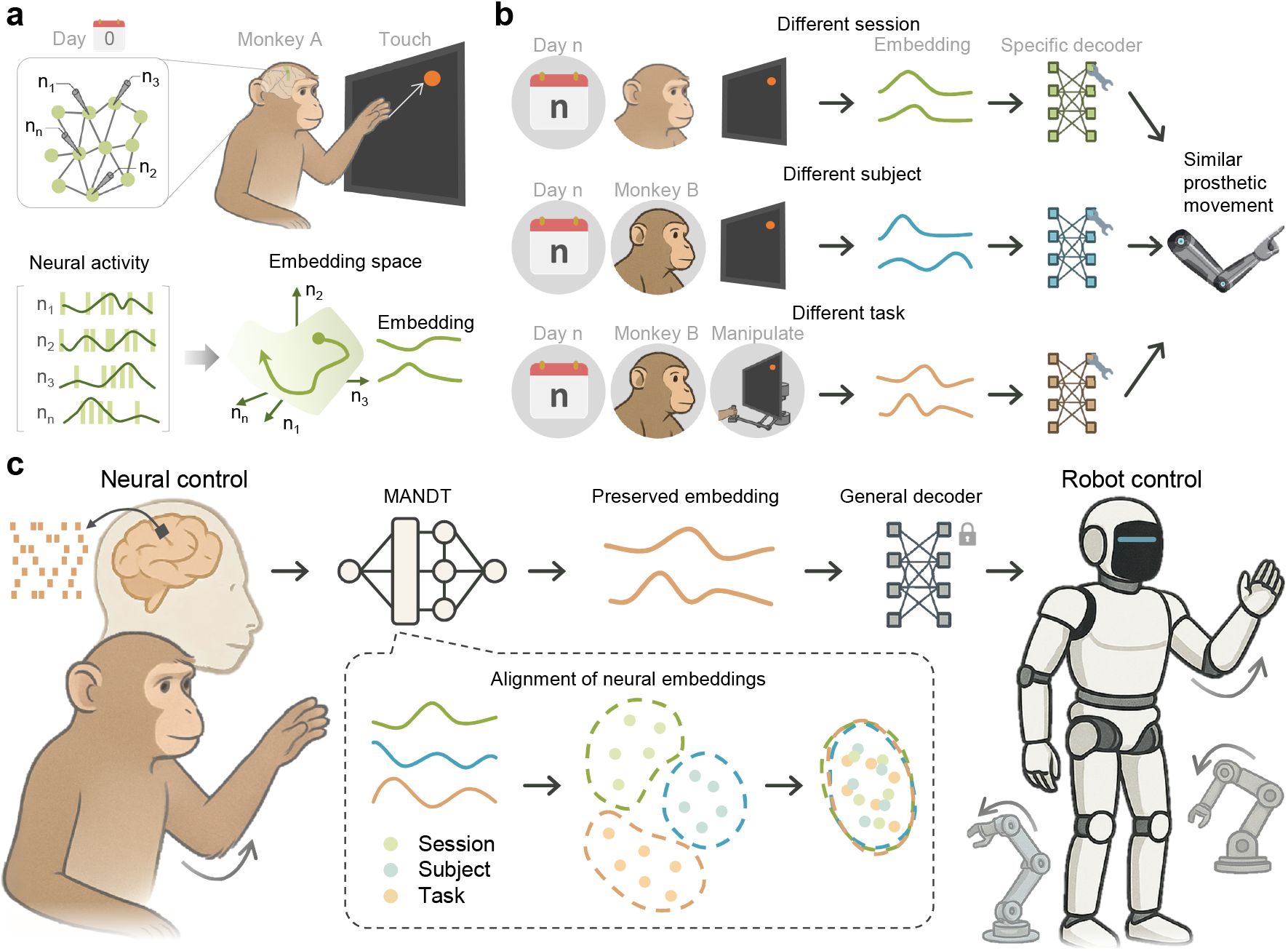
a, A monkey performs reach task while its neural activity in the motor cortex was recorded. Neural population activity can be described by low-dimensional dynamics embedded in N dimensions of neuronal activity. b, distinct neuronal activity in different sessions (days), subjects and tasks during similar behavior. c, From neural to humanoid control: MANDT (Multi-Aligned Neural Data Transformer) extract preserved embeddings of the neural activity by aligning them into a shared latent space. A deoder that fixed across multi-contexts (session, subject and task) decodes the preserved embeddings, enabling the monkey to directly control humanoid robots.

The generalization of BCI decoders would be greatly improved if they decode the preserved embeddings of neural population activity that are invarient across seesions, subjects and tasks. For across-session stability, linear alignment reveals stable latent dynamics in the motor cortex during consistent behavoir [11]. The stablity of cortical dynamics also improved the stablitiy of decoder performance throughout long timespans. For across-subject stability, neural population dynamics are also preserved across individuals performing similar behaviors, and enabled reliable decoding across different individuals [12]. For across-task stability, the dominant neural covariance patterns show similar structure across different motor tasks. The largest component of neural activity remains task-independent [13], revealing the neural basis of behavioral repertoire.

Current methods to extract the preserved latent dynamics rely on capturing and aligning embeddings from neural recordings [14]. Linear approaches typically apply dimensionality reduction methods such as Principal Component Analysis (PCA) [15, 16], Factor Analysis (FA) [17, 18] or GPFA [19] to capture low-dimensional latent dynamics of cortical activity, and align dynamics using linear transformations like Canonical Correlation Analysis (CCA) [11] or Procrustes Analysis [20]. However, these approaches assume neural activity lies in Euclidean space and may miss complex manifold structures [21–23]. Deep learning methods have emerged to address these limitations by capturing non-linear relationships. Adversarial networks can maintain decoder performance across sessions [24, 25], but are limited to pairwise session alignment. Variational auto-encoder (VAE) approaches with ‘stitching’ strategies share core architectures while using session-specific input/output matrices [26]. This method improved multi-session decoding but with but with slow inference speed. Recently, transformer-based methods [27, 28] such as Neural Data Transformer (NDT) leverage masked autoencoders [29] and context embeddings for multi-context learning. However, the preserved latent dynamics across session, subject and task have not been explicitly revealed and investigated within a unified computational framework.

Here, we proposed Multi-Aligned Neural Data Transformer (MANDT), a framework that preserved essential neural geometry while filtering out domain-specific noise. This framework reveals consistent embeddings of the motor cortex during similar movements across different sessions, subjects, and tasks. Our approach addresses a critical real-world challenge-unbalanced data distributions across experimental conditions for effective embeddings of cortical dynamics. Moreover, we decoded the extracted embeddings of one session data, achieving robust decoding performance across multi-contexts. We demonstrated the generalization of our framework across three distinct robotic platforms. Notably, we achieved direct neural control of a humanoid robot performing complex reaching movements, showing the intersection of biological and artificial embodied systems.

## 2 Results

### 2.1 Neural population dynamics modeling with transformer architecture

Neural population recordings exhibit substantial variability across sessions, subjects, and behavioral contexts, posing significant challenges for robust neural decoding applications. To address this challenge, we proposed a Multi-Aligned Neural Data Transformer (MANDT) that learns preserved embeddings of neural population activity.

Our approach is inspired by the masked language model (MLM) [30] that learns neural population dynamics by predicting masked spike counts based on surrounding context. The model operates in two modes: a single-session variant for within-session analysis and a multi-context variant for cross-context generalization. For the single-session model (Fig. 3a), spike trains were first embedded into a latent space by a linear layer, and incorporate spatial and temporal information. Then a spatiotemporal transformer block simultaneously processes spatial relationships and temporal dynamics of the latent states. After Gaussian smoothing, a bottleneck layer compresses high-dimensional signals into a reduced feature space, then reconstructs the activity through a residual connection. For the multi-context scenario (Fig. 3b), we incorporated learnable session-specific alignment modules that adapt embeddings across different recording sessions, and task-specific classification heads for movement direction prediction. Neural embeddingss are shuffled during training for robust learning of consistent embeddings under similar behavoirs regardless of recording session or trial.

**Fig. 3:**
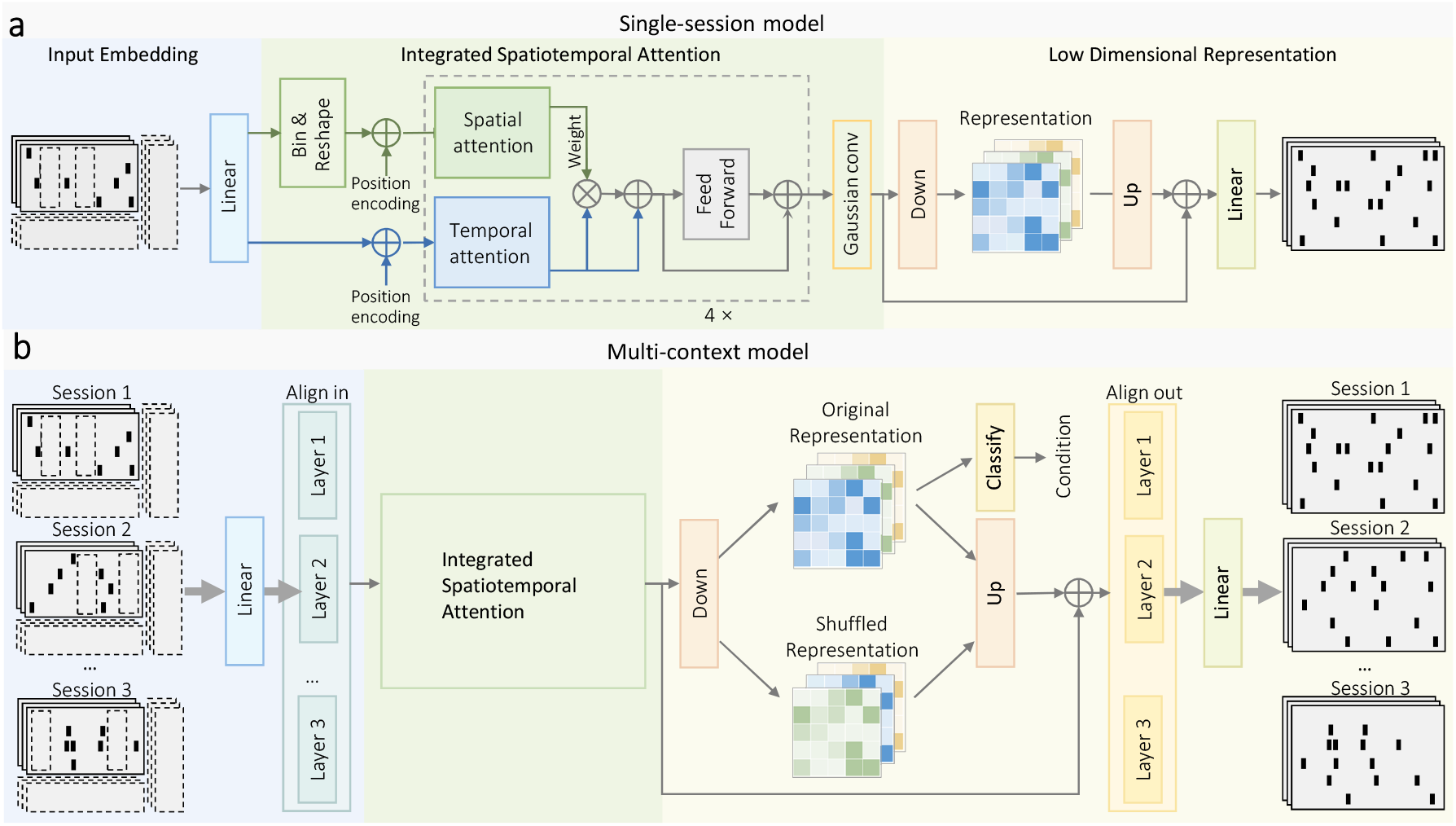
Model architectures. a, Single-session model that consists of a linear embedding layer that projects the neural activity into a latent space, a integrated spatiotemporal attention module to learn spatiotemporal features, a pair of ‘down’ and ‘up’ layers to extract low dimensional embedding, and a output layer to generate neural activity. b, Multi-context model for preserved embeddings extraction from different sessions, subjects and tasks. The model were based on the single-session model, but used alignment-in module to project neural activity into a shared latent space and a alignment-out module to map into specific latent space. We shuffled the embeddings to encourage the model to learn preserved embeddings across multi-contexts.

MANDT models single-trial spike counts at time *t* as Poisson-distributed observations generated from underlying firing rates [26]. Thus, the inferred firing rates are compared against the observed spike counts to compute negative log likelihood loss with Poisson distribution. The model is optimzed to predict the maksed spiking activity while maintaining sensitivity to behavioral variable.

### 2.2 Neural reconstruction across multi-contexts

To evaluate the generalizability of our neural encoder model, we systematically tested its reconstruction performance across increasingly challenging scenarios: single session, across different sessions, across different sessions and subjects, and across different sessions, subjects and tasks. This progressive evaluation framework allowed us to assess how well our model infers neural population dynamics while accommodating sources of variability in neural recordings. First, we analyzed motor cortical recordings from monkeys performing an instructed-delay reaching task, where two monkeys performed standard reaching task and a monkey performed maze reach task (Fig. 5a, b and Supplementary Fig. 1).

We established baseline performance by evaluating single-session model, which reconstructed firing rates with smaller trial-by-trial variability across repetitions of the same condition than real neuronal responses (Fig. 5c, first two columns), indicating effective denoising of spiking variability while preserving essential temporal structure. However, LFADS-inferred rates displayed smaller noise than that from our model. We evaluated reconstruction quality using two complementary metrics: co-bps (co-smoothing bits per spike), which measures how well the model predicts held-out neural activity [31], and all-bps (all-smoothing bits per spike), which assesses overall reconstruction ability across all recorded spikes. Our model achieved high reconstruction quality, with higher all-bps than SLDS [32], LFADS [26], and NDT [27], and higher co-bps than smoothing [31], GPFA [19], SLDS, and NDT (Fig. 5d). Note that our model selection was based on overall reconstruction quality (all-bps), which differs from methods (LFADS and NDT) that optimize for masked neural activity (co-bps).

Furthermore, we evaluated the multi-context model’s ability to generalize across sessions within the same subject. Our model reconstructed firing rates better than LFADS, but has higher trial-by-trial variability than that from single-session model (Fig. 5a, third column). Importantly, the reconstruction quality remained comparable to that of the corresponding single-session models (Fig. 5f and Supplementary Fig. 2), indicating that our model effectively captured the neural population activity across recording sessions.

We next evalated our model with cross-subject reconstruction, a more challenging task requiring accommodation of both temporal variations and inter-subject differences in neural anatomy. Our model maintained small trial-by-trial variability (Fig. 4a, fourth column). The cross-subject model achieved co-bps and all-bps performance comparable to the cross-session model for monkey C dataset, but showed an interesting trade-off pattern for monkey M dataset: reduced co-bps but increased all-bps compared to single-session performance (Fig. 4f and Supplementary Fig. 2). This precision trade-off reflects the model’s adaptation to accommodate temporal and inter-subject variability while maintaining overall reconstruction quality.

**Fig. 4:**
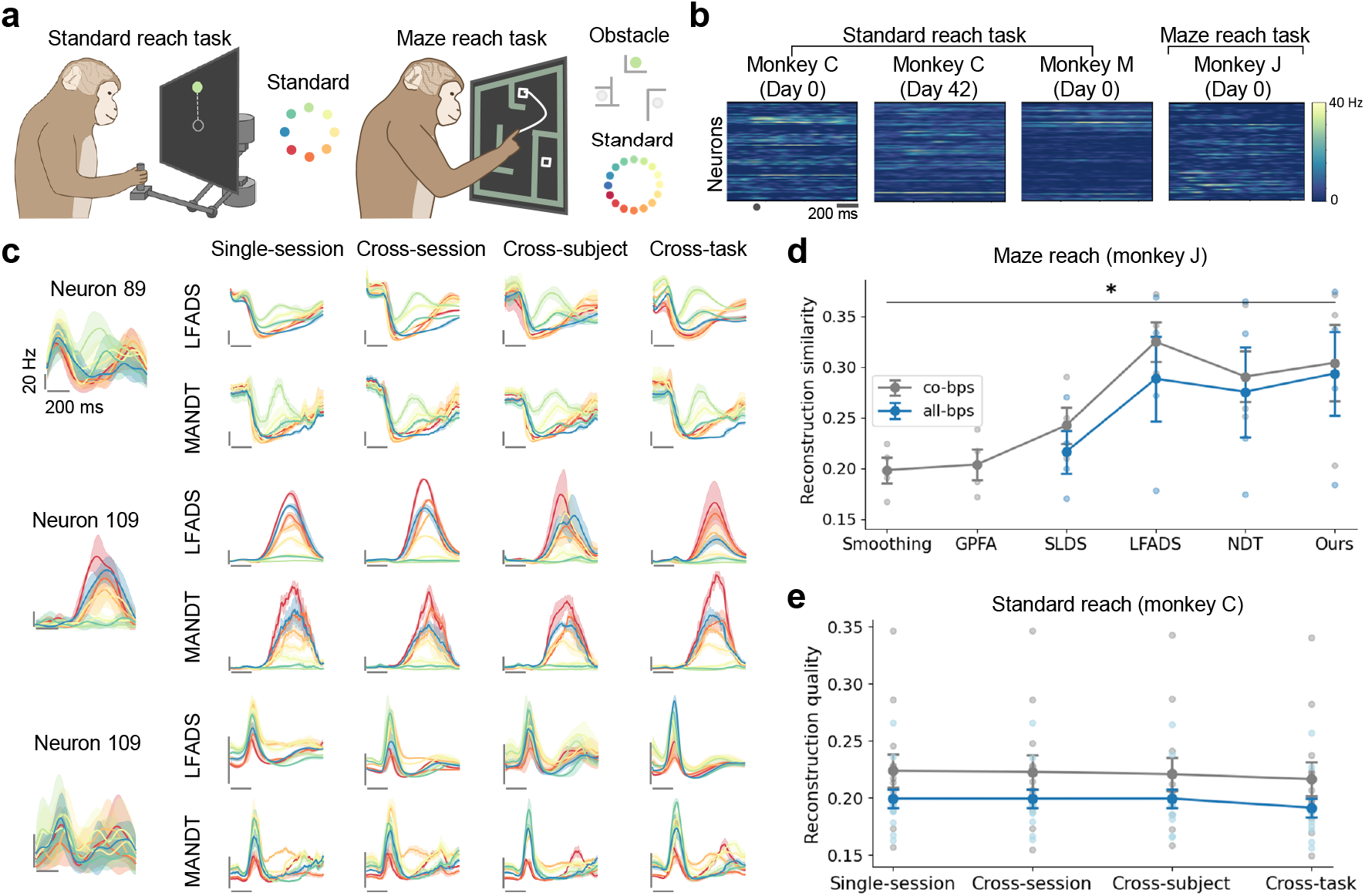
Reconstruction of neural activity. a, Experimental paradigm. Left, Standard reach task. Monkeys initiated each trial by placing their hand at the center of the workspace. A peripheral target appears after a random delay, and monkeys were required to wait until a go-cue signal before reaching toward the target. Right, Maze reach task where monkeys reach to the target while avoiding obstacles. b, Example neural firing rates aligned to movement onset for different recording session, subeject and task when monkey perform upward reaching. Circle denotes time of movement onset. c, Trial-averaged real and inferred rates for neurons for MANDT and LFADS within single session and across multi-context (session, subject and task). Color corresponds to each reach direction and shading indicates standard error. d, Neural reconstruction quality for monkey J dataset for different methods. e, Neural reconstruction quality for monkey C dataset within single session and across multi-context (session, subject and task).

Finally, we performed the most challenging test involved cross-task reconstruction, combining session, subject, and task variability. We trained our model on both standard reach tasks (monkeys C and M) and maze reach tasks (monkey J), hypothesizing that despite differences in movement planes and target distances, the underlying neural dynamics should share common latent structure. The predicted firing rates of the across-task model still maintain small trial-by-trial variability (Fig. 4c, last column). The reconstruction quality across-task are slightly smaller than that of the across-session and across-subject models (Fig. 4f and Supplementary Fig. 2). This performance degradation reflects the additional complexity of accommodating task-specific variations while preserving shared cortical dynamics.

In summary, our multi-context model can achieve stable reconstruction performance across multiple contexts (session, subject and task), overcoming the generalization challenges of neural plasticity, individual difference and task variations.

### 2.3 Preserved embeddings across multiple contexts

It is expected to observe consistant neural embeddings of similar behavior regardless of different contexts. We validated MANDT’s consistency across three progressively challenging scenarios: cross-sessions, cross-subjects, and cross-tasks.

For the cross-session consistency, the components of embeddings showed similar patterns across recording sessions while maintaining distinct signatures for different movement conditions (Supplementary Fig. 3a). We visualized the neural trajectory by using an orthonormalization procedure of the extrated embeddings (Fig. 5a, b, Supplementary Fig. 3a, b). MANDT neural trajectory displayed similar patterns within the same reach and circular configuration across reaches. LFADS [26] produced smooth trajectories but with different circular configuration. CEBRA-behavior [33] captured spatial information but lacked clear temporal structure. Aligned-PCA [33] showed tangled trajectories with high trial-to-trial variability despite preserving global reach arrangements.

**Fig. 5:**
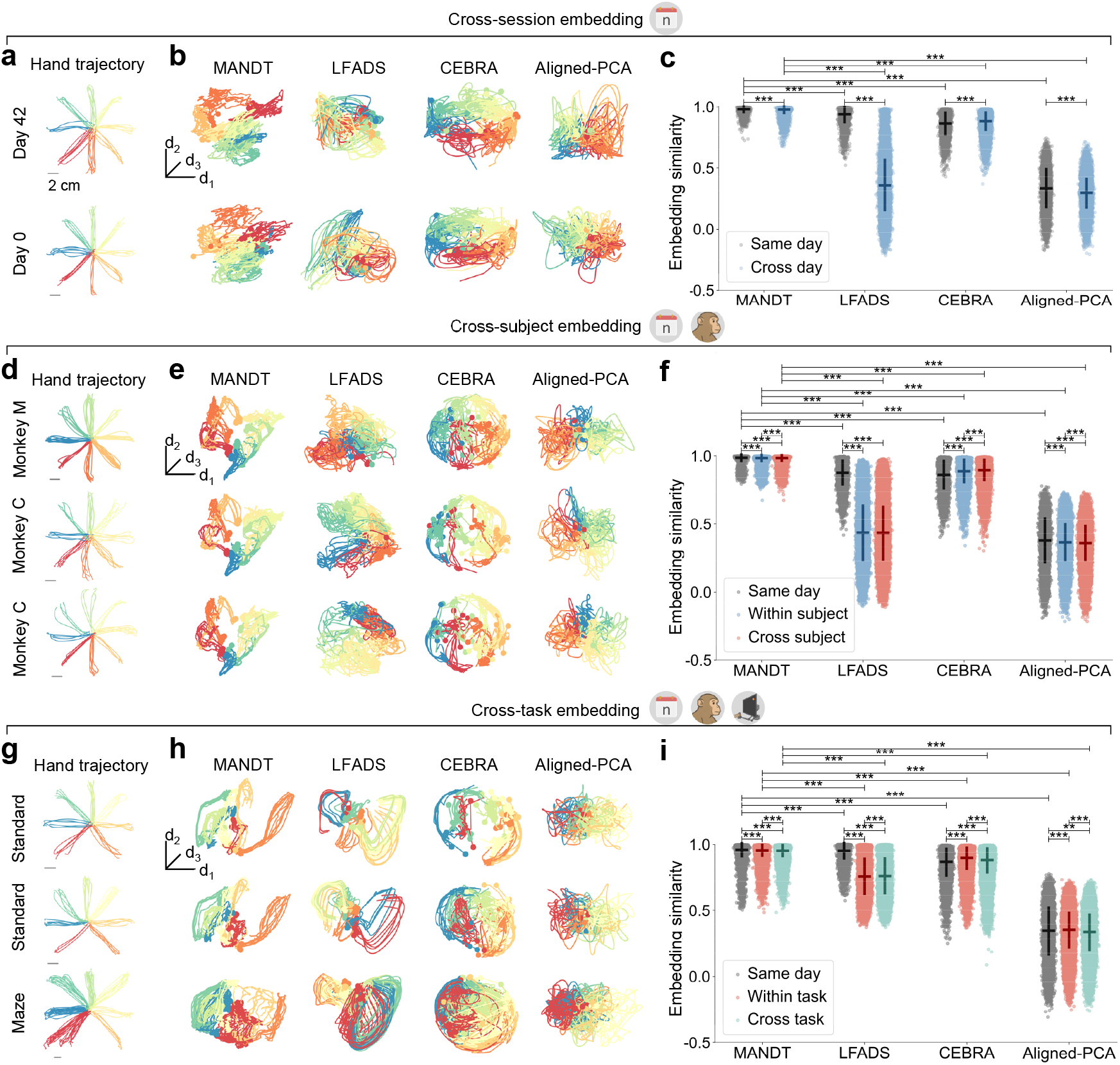
Consistent latent dynamics across multiple contexts. a, hand trajectory of two different sessions of monkey C. b, Orthonormalized neural trajectories for MANDT, LAFADS, CEBRA and aligned-PCA corresponding to different recording session. c, Cosine similarity of the embeddings of trials within in the same session (day) and across session (day). Two-sided Wilcoxon tests (paired samples), *P <* 10^−2^; *P <* 10^−3^. (d-f) Same as (a-c) but for different sessions and subjects (monkey C and M). (g-i) Same as (a-c) but for different sessions, subjects and tasks (standard and maze reach tasks).

Quantitative analysis demonstrated exceptional consistency: cosine similarity between embeddings of identical movement conditions remained high across all trial pairs (0.976 ± 0.008, monkey M), despite most pairs originating from different sessions (Fig. 5c and Supplementary Fig. 3b). By examining the distribution of cosine similarity values for trial pairs from the same session (same day) versus different sessions (cross day), the consistency across the same day is significantly higher than that of cross day. This consistency was maintained across all reaching directions (Supplementary Fig. 3c). The consistency of MANDT embeddings is significantly higher compared with LFADS, CEBRA-behavior and aligned-PCA (Supplementary Fig. 3c). Moreover, neural embeddings across sessions within monkey C (0.983 ± 0.005) or monkey J (0.950 ± 0.009) also showed high consistency (Supplementary Fig. 4). These results suggested that MANDT identified consistent latent dynamics across sessions.

Extending to cross-subject analysis (including diffenrent sessions and subjects). MANDT preserved global circular configurations and temporal neural trajectories (Fig. 5d, e). While embeddings showed distinct components reflecting individual differences, they maintained highly similar patterns (Supplementary Fig. 3d). This result may reflect the model’s adaptation to balance individual variability across subjects and sessions. By comparison, LFADS produced subject-specific embedding spaces with limited generalizability. CEBRA-behavior captured spatiotemporal structure but with less clear pattens than MANDT. Aligned-PCA preserved the spatial structure but with large noise (Fig. 5e).

Cross-subject embedding similarity remained remarkably high (0.987 ± 0.002) (Fig. 5f and Supplementary Fig. 3e). The cosine similarity values for pairs of trials from the same session (same day), the same subject (within-subject) and different subjects (cross-subject) became larger (Fig. 5f and Supplementary Fig. 3f). When using MANDT, the consistency of the embedding across subjects is significantly higher compared with LFADS, CEBRA and aligned-PCA. Hence, MANDT can extract global geometric information despite the increased complexity introduced by inter-subject variability.

For the cross-task analysis that involved simultaneous variation across sessions, subjects, and tasks. MANDT embeddings corresponding to similar behaviors still exhibited remarkable similarity (Fig. 5g, h and Supplementary Fig. 3g). MANDT could still unfold global geometric information within neural recordings. While LFADS maintained temporal consistency but lost spatial structure, and CEBRA-behavior revealed global reach arrangements with mixed neural states, aligned-PCA failed to extract meaningful structure (Fig. 5h). Besides, MANDT maintained high embedding similarity (0.955 ± 0.008) (Fig. 5i and 3h), significantly exceeding all comparison methods. The cosine similarity from the same task (within-task) was significantly higher than that of the across-task (Fig. 5i and Supplementary Fig. 3i), suggesting task-specific differences in neural dynamics.

Collectively, our model extracted stable neural dynamics that remain consistent across multiple contexts in neural recordings. The high degree of alignment observed across sessions, subjects, and tasks provides strong evidence for the existence of preserved computational principles in neural population dynamics during motor control.

### 2.4 Robust across-context decoding

We assessed the generalization performance of a behavior decoder trained on the extracted embeddings. A decoder trained to predict hand position from the embedding of one session was tested on data from different sessions, subjects, and tasks (Fig. 6a).

**Fig. 6:**
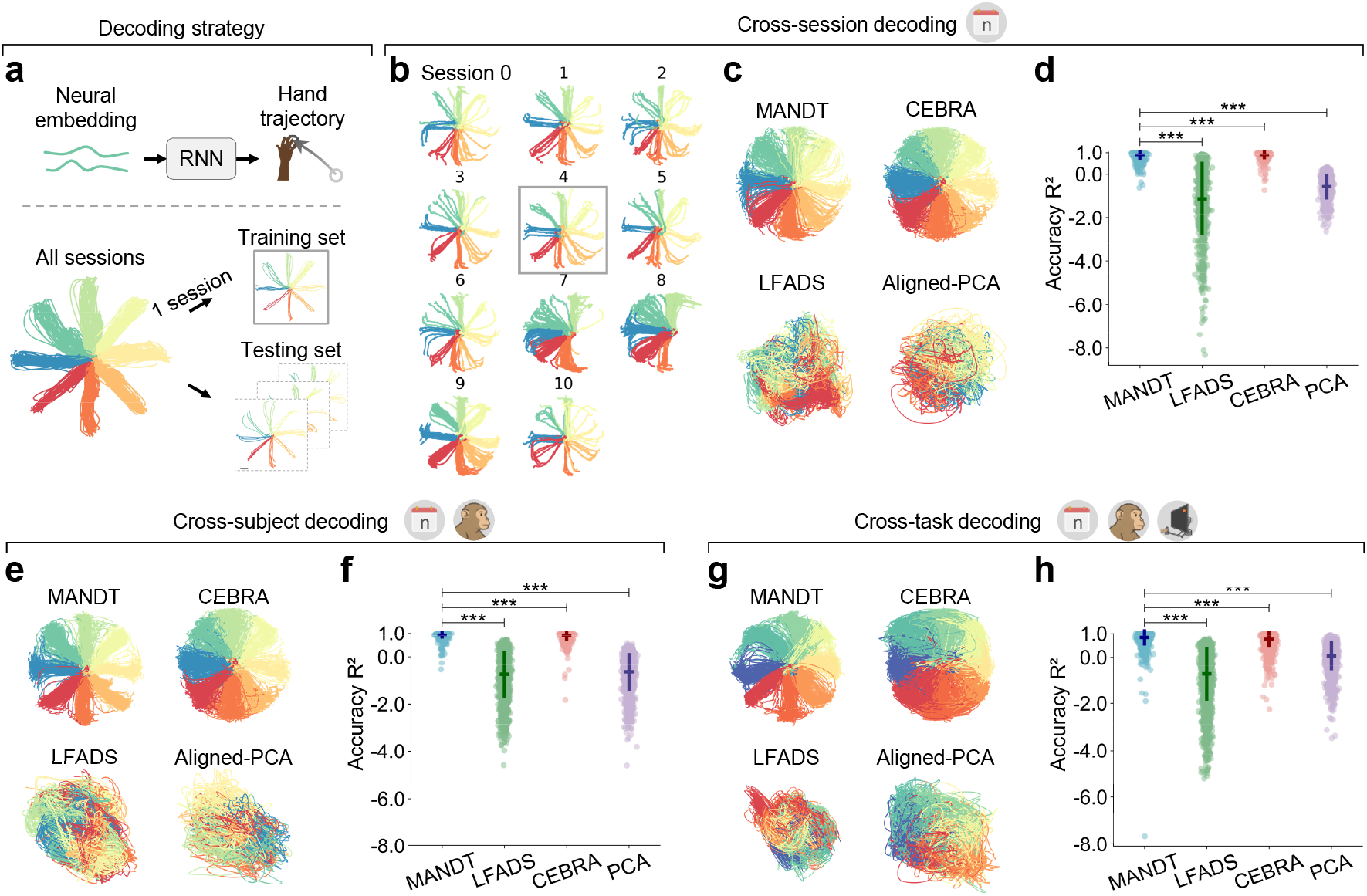
Decoding performance. a, Decoding strategy. Decoder (RNN) are trained to decode hand trajecty from neural embedding of one session data, and evaluated on the other sessions data. b, Decoded hand trajectories of each session within monkey M; the training session was boxed. c, Hand trajectories of monkey M decoded from embeddings of MANDT, LAFADS, CEBRA and aligned-PCA. d, *R*^2^ distribution of decoded hand trajectory using MANDT, LAFADS, CEBRA and aligned-PCA. Two-sided Wilcoxon tests (paired samples), *P <* 10^−3^. (e-f) Same as (c-d) except for cross-subject decoding. (g-h) Same as (c-d) except for cross-task decoding.

For the cross-session decoding generalization within the same subject, the behavior decoder that trained on the embeddings of one session of monkey C, decoded similar hand trajectories when applied to new sessions, with an *R*^2^ of 0.97 (Fig. 6b) and a mean absolute error (MAE) of 0.32 cm. Similar robust performance was observed across monkeys M (*R*^2^= 0.92, MAE = 0.57 cm) and monkey J (*R*^2^ = 0.96, MAE = 0.85 cm) (Supplementary Fig. 5d), confirming that our encoder extracts preserved neural dynamics across sessions. The decoded trajectories of MANDT were smoother than CEBRA-behavior (Supplementary Fig. 5a). However, decoding accuracy of MANDT at monkey C dataset was lower than those of LFADS and CEBRA-behavior, as some trials at the end of the movement are poorly decoded (Fig. 6c, d).

For the decoding generalization across sessions and subjects, when trained on one session of monkey C data and tested on both monkey C and M datasets, it maintained high accuracy on the both datasets (*R*^2^= 0.93 and MAE of 0.41 cm). Individual subject performance remained consistent: monkey C (*R*^2^ = 0.94, MAE = 0.37 cm) and monkey M (*R*^2^ = 0.93, MAE = 0.44 cm). The performances of the across-session and across-subject decoders are comparable with similar predicted hand trajectories (Supplementary Fig. 5d), demonstrating that our model captured common neural dynamics across subjects. The decoded hand trajectory showed correspondence to ground truth and substantially better than LFADS, CEBRA-behavior and aligned-PCA (Supplementary Fig. 5e, f). LFADS decoded wrong direction of some sessions due the instablity of embedding across subjects. The trajectory decoded from CEBRA dispeased within the same reah while displaying no clear structure from aligned-PCA (Supplementary Fig. 5b).

Cross-task generalization presented a greater challenge but remained robust. A decoder trained on the maze reach task achieved *R*^2^ of 0.93 and MAE of 0.74 cm across all contexts. However, the decoding performance degraded slightly compared to the across-session model (Fig. 6c, Supplementary Fig. 5a): *R*^2^ was decreased by 6.1% (0.97 to 0.91) on monkey C dataset, decreased by 6.5% (0.92 to 0.86) on monkey M dataset, but remained nearly identical for monkey J. The performance degradation may result from the involvement of curve reaches in monkey J’s maze task, which could disturb decoding straight reaches. Cross-session and cross-subject decoding performance yielded comparable *R*^2^ distributions, whereas cross-task performance exhibited significantly higher variability (Fig. 6d), with MAE distributions following a similar pattern (Supplementary Fig. 5b). MANDT outperformed competing methods significantly in behavoir decoding, maintaining comparable performance to cross-session decoding (Fig. 6g,h and Supplementary Fig. 5c). Overall, MANDT enables the decoder that trained on one session neural recording, maintain high decoding accuracy across all sessions. The stability of neural embedding allows robust decoding performance in BCI without recalibration.

### 2.5 Generality on cross-robot deploy

Teleoperation is a key application of the BCIs, requiring robust neural decoding that generalizes across diverse mechanical embodiments and task contexts. To demonstrate the broad applicability of our MANDT framework beyond laboratory settings, we systematically evaluated its performance across three distinct robotic platforms with increasing morphological complexity and degrees of freedom (DOF).

First, we deployed MANDT on a 6-DOF industrial robotic arm (JAKA, mini) for 2D trajectory reproduction, where the decoded hand position are fed to an inverse kinematics solver to compute joint angles for the path following (Fig. 7a). MAE between intended and executed paths was 1.2±0.2 mm. The robotic system maintained smooth, continuous motion with no observable discontinuities or jerky movements.

**Fig. 7:**
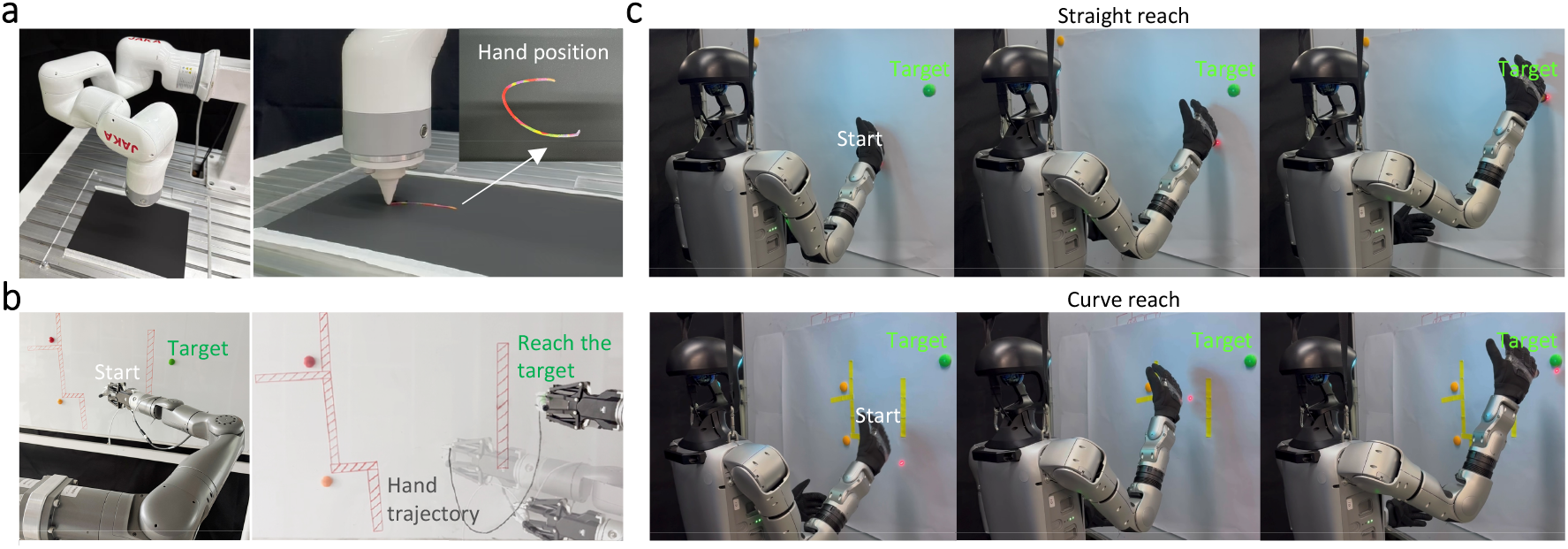
Time-series visualization of multi-platform robotic control. a, Industrial robotic arm validation. Robotic arm draws the decoded hand trajectory. b, Collaborative robotic arm validation. Robotic arm reaches to the target while avoiding barriers on the whiteboard. c, Humanoid Robot validation of the standard (top) and maze reach tasks (bottom). Multiple sub-images illustrate the execution process from movement onset to reaching the target.

Furthermore, we implemented 3D control using a 7-DOF collaborative arm (JAKA, K1) to investigate performance under kinematic redundancy. The increased DOF introduces an infinite solution space for inverse kinematics, requiring smooth and continous decoded path for robotic control. The robotic arm displayed smooth reaching behaviors and natural obstacle avoidance (Fig. 7b). However, MAE between intended and executed paths was 8.2 ± 2.1 mm, representing a performance decrease compared to the 2D drawing task due to higher-dimensional complexity.

Finally, we evaluated our framework on the humanoid robot platform to assess neural control when robotic and biological morphologies are highly similar. Humanoid robots closely approximates human arm anatomy and are capable of performing human-like tasks [34–36]. The humanoid robot held a laser pointer and pointed the decoded hand position on the whiteboard. The humanoid robot completed straight reach smoothly, with inter-joint coordination patterns similar to human movement (Fig. 7c, top). It also successfully executed curve reaching while avoiding barriers with natural fluidity (Fig. 7c, bottom). These results suggested that movement naturalness increased as the platform complexity increased.

In summary, we performed systematic validation across three robot s in real-world scenarios, progressively approaching human-like morphology and dynamics. Humanoid robots displayed most natural movement, making them promising in clinical BCIs for motor restoration.

### 2.6 Spatiotemporal attention mechanism reveals consistent neural processing patterns

To understand how our model processes neural population dynamics, we analyzed the attention mechanism across different contexts. First, we examined the spatial and temporal attention maps in the across-session model under one reach condition. The spatial attention maps exhibited sparse vertical patterns, indicating that a few dimensions are consistently attended to by all neurons. The attention map becomes smaller and denser as the model goes to deeper layers (Fig. 8a, left). On the other hand, temporal attention patterns of early layers displayed local temporal dependencies, with diagonal patterns reflecting attention to adjacent time points. Deeper layers exhibited complex patterns that deviated from the diagonal, suggesting that the model learned long-range dependencies in neural activity (Fig. 8a, right).

**Fig. 8:**
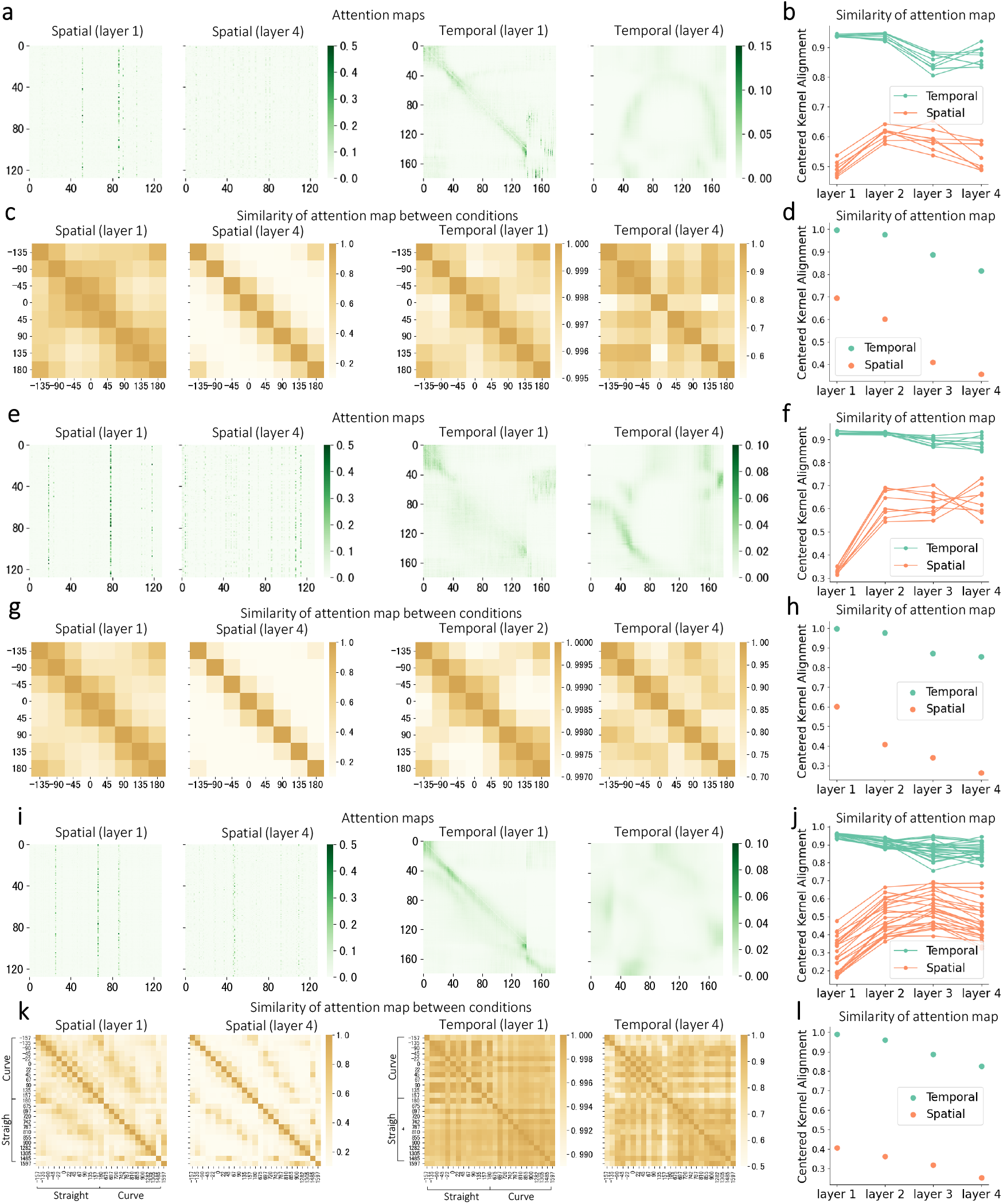
Spatiotemporal attention mechanism in inferring latent dynamics. a, Attention maps of spatial and temporal layers of across-session model. We visualized the first and last attention layers. b, Similarity of attention maps of across-session model between different trials (including different conditions). c, Similarity of attention maps of across-session model between conditions. d, Mean similarity value of (c). (e-h) Same as (a-d) except for across-subject model. (i-l) Same as (a-d) except for across-task model.

To quantify the consistency of attention mechanisms across trials, we computed similarity between attention layers using centered kernel alignment. Spatial attention maps showed low similarity in early layers but converged to higher similarity in deeper layers, while temporal attention maintained consistently high similarity across all layers, with slightly higher values in the early layers (Fig. 8B). This pattern suggests that the model employs a hierarchical processing strategy where early stages capture trial-to-trial variability while later stages extract consistent spatial representations. The high temporal similarity across layers suggests that movement temporal dynamics are relatively stereotyped across sessions.

We extended this analysis to examine attention patterns across different reach conditions. For spatial attention, both layers exhibit clear diagonal structure (Fig. 8C, left), indicating that similar movement directions share more similar attention patterns. For temporal attention, both layers display uniformly high similarity patterns across all condition pairs (Fig. 8c, right), revealing conserved temporal dynamics. Additionally, attention similarity between different reach conditions decreased progressively through deeper layers (Fig. 8d), reflecting the transformation from “noisy but rich” to “clean and discriminative” features through layered processing.

Furthermore, we extended our analysis to the attention mechanism in the across-subject model. The patterns remained consistent with the across-session model. Spatial attention maps displayed vertical patterns in early layers and became smaller at late layers, while temporal attention showed progression from local dependencies to complex long-range relationships (Fig. 8e). The similarity of attention layers between different trials within the same condition also replicated previous findings. Spatial attention similarity was lower than temporal attention similarity and increased at the late stage (Fig. 8f).

Cross-condition comparisons within the across-subject model revealed similar diagonal structures in spatial attention and uniform high similarity in temporal attention (Fig. 8g,h), confirming that the model utilizes consistent attention mechanisms for inferring neural population dynamics across subject and sessions.

Finally, we evaluated attention mechanisms in the across-task model, which encompassed increased complexity and diversity of behaviors. The fundamental attention patterns remained consistent with those observed in cross-session and cross-subject models. Spatial attention maps maintained the characteristic vertical structure, while temporal attention patterns showed similar local-to-global progression from early to late layers, though with increased complexity reflecting the varied temporal structures across different tasks (Fig. 8i, j).

The similarity analysis across reach conditions in the across-task model revealed task-specific modulation of attention patterns. Spatial attention similarity showed a block-diagonal structure, with higher similarity within the same task type and lower similarity across different tasks (Fig. 8k, left). This result suggests that while the model learns shared spatial representations, it also maintains task-specific spatial attention patterns. Temporal attention similarity were also higher across conditions and exhibited difference in straight and curve reaches (Fig. 8k, right), indicating that temporal dynamics of straight and curve reaches are slight different. The trend of attention map similarity was similar to that of the cross-session and cross-subject models (Fig. 8L).

In summary, our model utilizes a hierarchical attention mechanism that is consistent across sessions, subjects, and tasks. The spatial attention mechanism maintains information specificity at early stages and forms generalizable representations at later stages, while the temporal attention mechanism remained highly similar across stages. Our model also extracted the directional tuning property [37] of motor cortex data.

## 3 Discussion

The stability of the decoding performance of BCIs is still far from satisfactory due to the changes of extracellular environment [38], inter-subject anatomical variability [39], and task-specific neural dynamics [13]. Here we proposed a computational framework that learns preserved embeddings of neural recordings despite these three major sources of variability. The preserved embeddings allow decoders maintain robust performance when applied to new recording sessions, subjects and tasks, enabling zero-shot generalization. Moreover, we demonstrate the robustness of our computational framework across multiple robotic platforms, especially in humanoid robots, validating our framework’s adaptability to various real-world scenarios.

We leverage the aligned low-dimensional dynamics in neural population activity, which captures fundamental features of cortical computation [21]. Although recorded neurons display distinct patterns, their population activity are confined to a low-dimensional manifold spanned by specific patterns of correlated neural activity [21]. Studies have shown the existence of a low-dimensional neural manifold that remains consistent across time when aligning latent dynamics, suggesting that the embeddings of motor command within this manifold exhibits long stability [13, 26]. Moreover, recent evidences support the shared latent dynamics across individuals using manifold alignment [12, 33, 40]. Our framework exploits this principle by embedding the neural activity into a latent space and aligning latent dynamics by learnable session-specific transformations. Our method is consistent with the findings that motor command is encoded in stable, low-dimensional latent variables rather than in the activity of specific neurons [11, 12, 41, 42].

A critical architectural consideration in modeling concerns whether to use separated spatiotemporal attention (as in STNDT [43]) versus unified patch-based attention (as in NDT2 [28]). Unlike image patches where adjacent pixels typically belong to the same object or texture, neurons within a spatial “patch” may exhibit heterogeneous firing rates. In addition, neural correlations are not necessarily determined by physical electrode proximity—two spatially distant neurons may be highly correlated in their responses to specific behaviors, while adjacent neurons may serve different computational roles. Therefore, we used separated attention mechanism with integrated spatiotemporal processing. Our architecture integrates spatiotemporal processing within each transformer layer rather than separate parallel streams in STNDT [43]. The optimal architectural choice likely depends on the primary objective: separated attention appears more suitable for scientific understanding of neural circuit organization and interpretable decoding, while unified attention may excel in engineering applications requiring robust performance across diverse recording conditions. Future work might explore hybrid approaches that combine learned functional groupings with separated attention mechanisms, potentially capturing the interpretability advantages of STNDT while maintaining the scalability and generalization capabilities of NDT2.

The demonstration of our framework across multiple robotic platforms suggests the balance between abstraction and specificity of motor cortical activity. The motor cortex appears to encode motor command in an abstract format across biological and artificial systems, yet specific enough to preserve spatiotemporal dynamics of a body. Moreover, the translation of neural signals from non-human primates to humanoid robot movements presents a novel experiment paradigm that combines BCIs and embodied artificial intelligence (embodied AI), representing a paradigm shift toward more naturalistic and intuitive human-machine interfaces [44]. Our framework provides a foundation for developing new embodied AI systems that can leverage the sophisticated sensorimotor integration capabilities of biological neural networks while operating through artificial robotic systems with high precision, strength and efficency. This hybrid approach may lead to embodied intelligence systems that surpass the capabilities of either biological or artificial systems alone.

## 4 Methods

### 4.1 Dataset Collection

We collected a dataset of motor cortical recordings from hesus monkeys performing center-out reaching tasks, based on previously published datasets[11, 45]. The dataset included neural activity recorded from the primary motor cortex (M1) and dorsal premotor cortex (PMd) across 3 rhesus monkeys, 27 sessions, and 2 task variants.

#### 4.1.1 Subjects and Neural Recordings

Neural data were collected from three adult rhesus macaques (Monkey C, M and J) implanted with Utah arrays (Blackrock Microsystems, 96 channels) in M1 and PMd. Single-unit and multi-unit activities were isolated by spike sorting techniques. Neural data were binned in 5 ms intervals and aligned to movement onset, defined as the time when hand velocity exceeded 5% of peak velocity. Following the Neural Latents Benchmark (NLB) protocol [31], we partitioned neurons into “held-in” and “held-out” populations. “Heldin” neurons had their recorded activities preserved while “held-out” neurons were not visible to the model and masked by zeros. We randomly selected “held-out” neurons from electrodes that recorded multiple neurons, maintaining a 3:1 ratio of “held-in” to “held-out” neurons. We analyzed data from 250 ms before to 650 ms after movement onset. To create a prediction task, we masked neural activities from 500-700 ms post-onset with zeros [31]. For multi-context model training, neural data were zero-padded to match the maximum number of neurons across all sessions.

#### 4.1.2 Behavioral Tasks

Monkeys performed center-out reaching tasks in which they controlled a cursor displayed on a monitor by touching the screen or moving a manipulandum. Each trial began with the cursor at a central position, followed by the appearance of a peripheral target. After a randomized delay (400-1000 ms), a go cue (e.g., the central target disappears) prompts the monkey to reach the target. The animal then moved the cursor to the target and finished the trial. We collected data from two task variants:

##### Standard reach task

Monkeys controlled a manipulandum in a 2D plane to reach a target in one of 8 equally distributed around a circle. There were 8 conditions corresponding to target direction. Each subject contains about 18 sessions, with each session consisting of approximately 150 trials and 200 neurons. The details of the standard reach task dataset is in table 1.

##### Maze reach task

This task consists straight and curve types according to the appearance of barrier. In straight type, similar to the standard reach task but with 9 or 36 types of target positions. In curve type, there were virtual obstacles on the screen and some conditions included distractor targets. Monkeys had to determine which target was reachable and touched the screen to reach a target while avoiding intervening barriers. Here we redefined the movement conditions based on the target angle and the presence of obstacles, where some targets with very close angles are classified as one condition. The details of the standard reach task dataset is in table 2.

#### 4.1.3 Data Augmentation

We employed a data augmentation technique based on temporal jittering, which introduce controlled variability in the timing of spike events. Previous studies have shown spiking variability where precise spike timing can vary due to biological noise or measurement artifacts. By randomly shifting spike times within a bounded range while preserving their overall structure, temporal jittering simulates spiking variability in neural responses, thereby improving the robustness of models during training.

We first generated a probabilistic mask to identify spikes eligible for jittering. Only spikes with nonzero counts and probabilities below jittering probability (*p*_*j*_ = 0.6) were selected for jittering. For each selected spike, a random temporal shift is sampled from a uniform distribution over [−4, 4]. The shifted spikes are relocated to their new temporal positions. To handle overlapping shifts, spike counts are updated using a scatter-add operation to ensure correct accumulation. The final augmented dataset is constrained to a range of [0, max spikes], where max spikes] is the maximum spike counts in the dataset.

#### 4.1.4 Data Partitioning and Sampling Strategy

To account for different condition definitions across the two reaching tasks, we reclassified the conditions based on two factors: (1) the angular position of the target relative to the central point, divided into 16 equally spaced conditions at 22.5° intervals (ranging from -180° to 180°), and (2) the presence or absence of barrier, resulting in a total of 16 × 2 = 32 conditions. However, within the barrier conditions, there were distinct configurations of targets and barriers that produced different behavioral patterns. To capture these differences, we further subdivided the conditions based on the similarity of trial-averaged hand trajectories. For embedding shuffling (described in 4.2.2), we used these refined condition labels: target angles plus 720° or 1440° for barrier conditions. This labeling scheme distinguishes between barrier configurations while preserving the angular periodicity for both condition types.

We sorted data points by their condition labels and required that each batch contains the neural data from the same condition (which could span across session, subject and task). For the training set, we employed an oversampling strategy to address the class imbalance: we randomly resampled from the available data to fill each batch for conditions with insufficient samples, which promoted balanced learning across all conditions. For the test set, we also grouped data by condition but sequentially sampled from each condition until all data were exhausted, preserving the instinct distribution of the test data. Both sampling strategies shuffled the data within each condition to ensure randomness and prevent order-related biases.

### 4.2 Model Design

Our goal is to learn preserved embeddings of neural population activity for downstream tasks such as behavioral decoding. We consider two scenarios: (1) single-session modeling where data comes from one recording session, and (2) multi-context modeling where we must handle variability across multi-context, including different recording sessions, subjects and tasks.

#### 4.2.1 Single-session Model

Our single-session model introduces a transformer architecture that processes neural data in both temporal and spatial dimensions simultaneously. Our model integrates spatiotemporal processing within each transformer layer as neural population dynamics exhibit intrinsic coupling between spatial and temporal patterns.

##### 1. Input Embedding

Neural data was embedded using a linear embedding and a learnable positional encoding to obtain **X** ∈ ℝ^*T ×*128^ (T is the number of time points).

##### 2. Integrated Spatiotemporal Attention

Each transformer layer first processes temporal dependencies using standard multi-head self-attention:

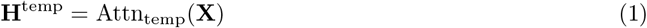

To capture population-level spatial interactions, we introduce a spatial attention mechanism. The temporal sequence is downsampled by averaging every 5 consecutive time steps to create spatial representations **X**^spatial^ ∈ ℝ^*T/*5*×*128^, which are then processed by spatial multi-head attention:

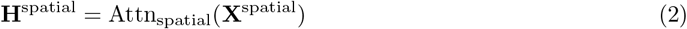

The spatial attention weights are then applied to the original temporal representations:

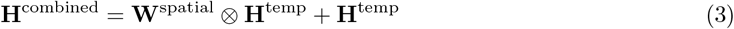

where **W**^spatial^ are the learned spatial attention weights. This integration allows spatial patterns to directly modulate temporal representations within each layer, rather than processing them independently as in STNDT[43].

To capture the prominent temporal structure, we applied Gaussian kernel (kernel size=4) to smooth the transformer output and interpolated it to the original time steps.

##### 3. Low Dimensional Embeddings

To encourage compact embeddings, we include a ‘down’ layer that projects the transformer output to a lower-dimensional space and an ‘up layer’ that projects to the embeddings to high-dimensional space:

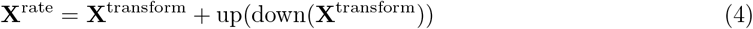

where down and up are ‘down’ and ‘up’ layers, respectively.

Finally, a linear output layer maps the **X**^rate^ into neural activity.

#### 4.2.2 Multi-context Model

Neural recordings often exhibit large variability across sessions, subjects and tasks. We extend the single-session model with session-specific transformations to handle this variability. The multi-session model is characterized with shared transformer parameters and session-specific alignment transformations. This allows the model to learn universal and robust embeddings patterns and adapt to session-specific variations.

##### 1. Alignment-in Module

Each session *s* has a learnable layer to transform the neural activity from different session into a common latent space:

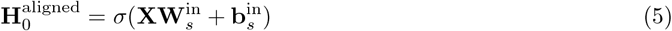

where **X** is the data after Input Embedding. 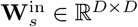 and 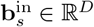 are session-specific parameters, initialized as identity transformations and zero biases respectively. *σ* is GELU function.

##### 2. Embedding Shuffling

To enforce that similar behaviors produce consistent embeddings regardless of recording session, we implemented a embedding shuffling strategy during training. After obtaining the initial output embeddings from the Integrated Spatiotemporal Attention block, we randomly shuffled the embeddings within a batch, creating a shuffled version. Both original and shuffled embeddings are then up-projected and combined with the rate output:

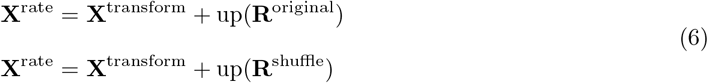

where **R**^original^ and **R**^original^ are the original and shuffled embeddings.

The original embeddings passed through a condition layer to predict movement conditions.

##### 3. Alignment-out Module

output embeddings are transformed by a session-specific layer to session-specific latent space:

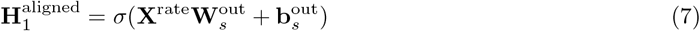

where 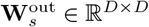 and 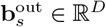 are session-specific parameters, initialized as identity transformations and zero biases respectively.

The shuffled output embeddings were also transformed through Alignment-out Module.

#### 4.2.3 Behavior Decoder

We trained a behavior decoder to map the extracted low-dimensional neural dynamics to hand position for robotic arm control. The behavior decoder consists of an RNN followed by a linear output layer. The RNN has 32 hidden units. The behavior decoder was trained to minimize the mean squared error between predicted and hand position:

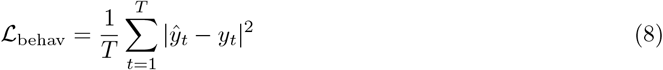

where *ŷ*_*t*_ is the predicted hand position, *y*_*t*_ is the actual hand position, and *T* is the number of time steps.

### 4.3 Training Objective

#### 4.3.1 Single-session Model

We adapted the masked language modeling objective from natural language processing to computational neuroscience. We masked held-out neurons, future activity and a fraction of time steps [27] and trained the model to predict the masked values. The loss function depends on the hypothesis of spike counts as Poisson-distributed, defined as:

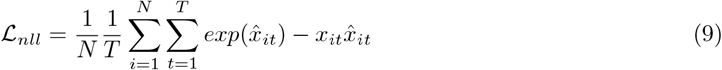

where *N* is the number of neurons and *T* is the number of timesteps. *x*_*ij*_ and 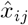 are the predicted firing rate and spike counts at neuron *i* and timestep *t* that are masked.

#### 4.3.2 Multi-context Model

The total loss combines neural reconstruction, preserved embeddings extraction and auxiliary condition classification:

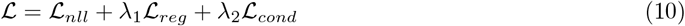

where *λ*_1_ = 5 × 10^−2^, *λ*_2_ = 5 × 10^−3^.

Under the hypothesis that neural dynamics in individuals are preserved when they perform similar behavior, we expect that the embeddings should be robust to cross-trial permutations underlying consistent behavior. Specifically, if multiple trials correspond to the same condition, the neural dynamics should follow similar trajectories in the latent space, regardless of which specific neural population instances are observed. Therefore, the regularization loss is formulated as:

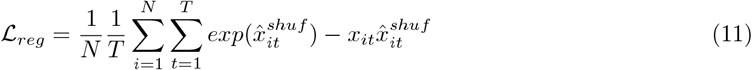

where 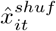 represents predictions from shuffled neural data.

For the auxiliary condition prediction task, we design a novel cosine embedding loss that leverages the periodic nature of angular data. Given discrete angle classes *θ*, we map predictions and ground truth to a 2D embedding space. The angle embedding function is defined as:

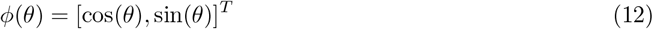

We compute a weighted combination of all possible angle embeddings:

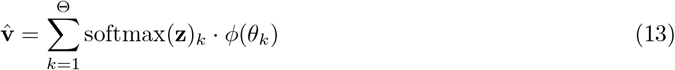

where **z** represents the prediction logits and *θ*_*k*_ ∈ Θ are the discrete angle values.

The ground truth embedding is computed directly:

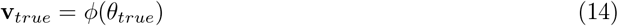

The cosine embedding loss is then formulated as:

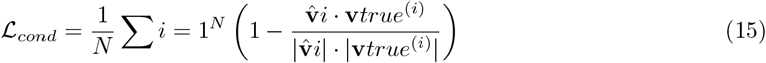

We used Adam with cosine learning rate scheduling and Bayesian optimization for hyperparameters tuning for both the models.

### 4.4 Evaluation

We evaluated the performance of of our framework in the aspects of neural activity reconstruction, latent dynamics similarity and kinematic decoding performance.

For neural reconstruction accuracy, we used co-smoothing (co-bps) [31], which measures the model’s ability to predict unseen (held-out) neural activity, defined as:

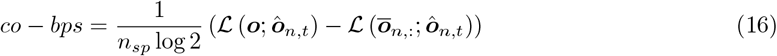

where ℒ is the sum of the log-likelihoods with Poisson distribution, ***o*** and ***ô*** are ground truth and predicted firing rates of held-out neurons, 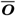 is the mean firing rate for the neuron *n* and *n*_*sp*_ is its total number of spikes. The larger the co-bps value, the better the model’s predictive ability.

We defined all-bps, measuring the ability of the model to reconstruct the complete neural activity. It is computed in the similar manner as co-bps but on the time points of all neurons.

For latent dynamics similarity, we used cosine similarity of the embeddings under similar behavior:

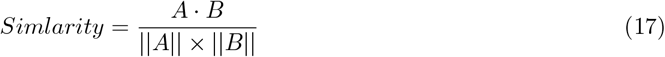

where *A* and *B* are flattened embeddings from different trials (sessions, subjects or tasks) but with similar behavior.

We also compared the similarity between attention map using centered kernel alignment (CKA), which is used to analyze the similarity between neural network layers or learned embeddings [46]. Given two kernel matrices *K* and *L*, CKA is computed as:

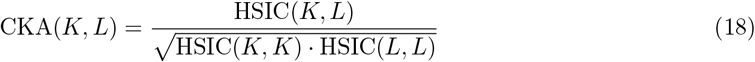

where: *HSIC* is a kernel-based independence measure:

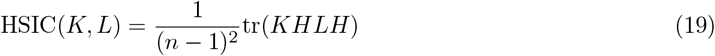

where 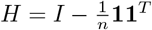 is the centering matrix (**1** is a vector of ones).

For behavior decoding accuracy: We computed the Mean Absolute Error (MAE), Root Mean Squared Error (RMSE) and Coefficient of Determination (*R*^2^) between actual and predicted hand position, defined as:

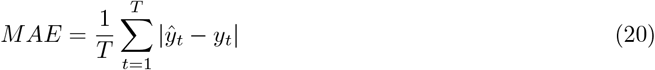

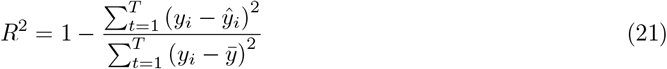

where ***y*** and ***ŷ*** are true and predicted hand positions, 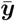 is the mean of ***y***.

### 4.5 Robotic Platforms

We evaluated the practical application of our framework using three robotic platforms with increasing complexity: industrial, collaborative, and humanoid robots. This hierarchical validation allowed us to assess our framework’s adaptability to various real-world scenarios. Neural signals were processed through our pretrained encoder-decoder model to generate hand trajectory predictions at 200 Hz. For each decoded position, the robot system calculated optimal joint configurations through inverse kinematics solution and applied joint control commands of the right hand. The cartesian poses of the rx, ry, rz were fixed during movement.

First, we validated on a 6-DOF industrial arm (JAKA, Mini 2), equipped with a pen holder, which executed decoded trajectories on paper within a calibrated 20×30 cm workspace. Second, we controlled a 7-DOF robotic arm (JAKA, K-1) to draw the decoded trajectory on a vertical whiteboard. Third, a humanoid robot (Unitree, G1) with 29 controllable degrees of freedom (38 DOF in total, joints below the waist were fixed) held a laser pointer and point the decoded hand position on a vertical whiteboard.

To accommodate different workspace constraints and mechanical properties, drawing trajectories were scaled down by a factor while screen control trajectories were scaled up by a factor of 2 to utilize the display area.

## Acknowledgments

This work was funded by the STI 2030—Major Projects (2022ZD0208604), and the National Natural Science Foundation of China (61773259). We are grateful to Student Innovation Center for platform support, to Prof. Xinyu Chai, Prof. Liming Li and Prof. Xiaohong Sui for instruction, to Jieji Ren for discussions.

## Code availability

The codes used in this study are available on Github at

## Author contribution

H.J. conceived the idea and designed the study. H.J., Z.L, X.B, A.L, X.J conducted the experiments. H.J. and Z.L analyzed and interpreted the results. Y.C directed the project. All the authors contributed to the writing and editing.

